# Interbacterial transfer of carbapenem resistance and large antibiotic resistance islands by natural transformation in pathogenic *Acinetobacter*

**DOI:** 10.1101/2021.08.05.455225

**Authors:** Anne-Sophie Godeux, Elin Sveldhom, Samuel Barreto, Anaïs Potron, Samuel Venner, Xavier Charpentier, Maria-Halima Laaberki

## Abstract

*Acinetobacter baumannii* infection poses a major health threat with recurrent treatment failure due to antibiotic resistance, notably to carbapenems. While genomic analyses of clinical strains indicate that homologous recombination plays a major role in the acquisition of antibiotic resistance genes, the underlying mechanisms of horizontal gene transfer often remain speculative. Our understanding of the acquisition of antibiotic resistance is hampered by the lack of experimental systems able to reproduce genomic observations. We here report the detection of recombination events occurring spontaneously in mixed bacterial populations and which can result in the acquisition of resistance to carbapenems. We show that natural transformation is the main driver of intra-, but also inter-strain recombination events between *A. baumannii* clinical isolates and pathogenic species of *Acinetobacter*. We observed that interbacterial natural transformation in mixed populations is more efficient at promoting the acquisition of large resistance islands (AbaR4, AbaR1) than when the same bacteria are supplied with large amounts of purified genomic DNA. Importantly, analysis of the genomes of the recombinant progeny revealed large recombination tracts (from 13 to 123 kb) similar to those observed in the genome of clinical isolates. Moreover, we highlight that transforming DNA availability is a key determinant of the rate of recombinants and results from both spontaneous release and interbacterial predatory behavior. In the light of our results, natural transformation should be considered as a leading mechanism of genome recombination and horizontal gene transfer of antibiotic resistance genes in *Acinetobacter baumannii*.

**Importance:** *Acinetobacter baumannii* is a multidrug resistant pathogen responsible for difficult-to-treat hospital-acquired infections. Understanding the mechanisms leading to the emergence of the multi-drug resistance in this pathogen is today crucial. Horizontal gene transfer is assumed to largely contribute to this multidrug resistance. However, in *A. baumannii*, the mechanisms leading to genome recombination and the horizontal transfer of resistance genes are poorly understood. We bring experimental evidence that natural transformation, a horizontal gene transfer mechanism recently highlighted in *A. baumannii*, allows the highly efficient interbacterial transfer of genetic elements carrying resistance to last line antibiotic carbapenems. Importantly, we demonstrated that natural transformation, occurring in mixed populations of *Acinetobacter*, enables the transfer of large resistance island mobilizing multiple resistance genes.

## Introduction

*Acinetobacter baumannii* is a gram-negative bacterium responsible for a wide range of infections in both humans and animals (1, 2). This multidrug-resistant (MDR) agent poses a health threat, particularly in intensive care units where it can lead to bacteremia and ventilator-associated pneumonia. Consequently, secondary infections with multidrug-resistant *A. baumannii* have been reported during the COVID-19 pandemic (3, 4). *A. baumannii* infections are steadily resistant to multiple antibiotics including to carbapenems. For Europe only, a combined resistance to fluoroquinolones, aminoglycosides and carbapenems is observed for nearly 30% of invasive *Acinetobacter* sp. isolates (5). In *A. baumannii*, carbapenem resistance is mainly associated with the expression of OXA23 carbapenemase encoded by the *bla*_OXA-23_ gene (6). The composite transposon Tn*2006*, formed by two ISAba1 insertion sequences framing the *bla*_OXA-23_ gene, is the main genetic context for the *bla*_*OXA-23*_ gene and can be found in different locations, including plasmids, but it is most often found inserted in the large resistance island AbaR4 (6, 7). Such large and diverse *A. baumannii* resistance islands (Ab-RI) are potential contributors to the multi-drug resistance phenotype observed in *A. baumannii*. The first description of an Ab-RI was reported in 2006 in the epidemic strain AYE with the AbaR1 island consisting in an 86 kb-long genomic structure (8). Since then, analysis of more than 3,000 *A. baumannii* genomes revealed that Ab-RIs are present in nearly 65% of them (9). These genomic island present a great diversity in gene content with often multiple putative antibiotic-resistance genes (10, 11). Ab-RI were presumed to be initially acquired through plasmid conjugation followed by chromosomal insertion, and evolve through multiple insertions and rearrangements of insertion sequences (12, 13). However, genome analyses also support their horizontal transfer between distantly related isolates, as exemplified by the acquisition of the 21 kb-long ABGRI3 resistance island (14) or the 35 kb-long AbGRI5 (15). Both acquisition involved large recombination events (up to 34 kb-long) at the sequence flanking the island. Indeed, high rates of genome recombination is a hallmark of *A. baumannii* genomes (16, 17). Recombination events may have led to the acquisition by MDR strains of *parC* or *gyrA* alleles conferring resistance to fluoroquinolones and of IS*Aba1* upstream of the *ampC* gene leading to resistance to 3^rd^ generation cephalosporins (14, 17, 18). Genome analysis of isolates from a longitudinal study also offered a glimpse of the gene transfer and recombination going on in the hospital setting, with the acquisition of the *bla*_OXA-72_ variant of the *bla*_OXA24/40_-type carbapenemase gene and recombination affecting the *bla*_*OXA-51*_ locus (19).

Despite their importance in the evolution of *A. baumannii* into an MDR pathogen, the mechanisms leading to genome recombination and recombination-dependent acquisition of resistance genes and Ab-RI remain elusive. Indeed, few studies have experimentally investigated the horizontal transfer of chromosomal antibiotic resistance genes in *A. baumannii*. The chromosomal transfer of an 11 kb-long Tn*215* harboring the *bla*_NDM-1_ during mating of the isolate R2090 to the reference strain CIP 70.10 was experimentally observed and involved the acquisition of a 65 kb region through homologous recombination (20). While transduction by prophages was suggested, the mechanism of transfer was not elucidated. Phage particles, present in prepared fractions from culture supernatant of the clinical isolate NU-60 were found to mediate chromosomal transfer to the reference strain ATCC17978 (21). If generalized transduction by strain-specific prophages is a potential mechanism of resistance gene transmission, natural transformation is another potential and highly conserved route. Natural transformation allows bacteria to actively import exogenous DNA and, provided a sufficient sequence identity, it integrates the recipient cell’s genome by homologous recombination (22). Most *A. baumannii* isolates and also closely related *Acinetobacter nosocomialis* were found capable of natural transformation when presented with purified DNA (23–27). Yet, we currently have a limited understanding of the role of natural transformation in genome dynamics and the spread of antibiotic resistance in *A. baumannii* populations.

In the current study, we explored the inter-species and inter-genus transfer of antibiotic resistance, notably to carbapenems, which is spontaneously occurring in mixed populations of pathogenic *Acinetobacter* sp.. We provide evidence that natural transformation is the main transfer route and fosters recombination events and the acquisition of multiple resistance genes carried by large genomic island. Experimentally replicating the large recombination events observed in *A. baumannii* chromosomes, our results suggest a major role played by natural transformation in the dynamics of *A. baumannii* genomes.

## Results

### Intra and interspecies recombinants are produced in mixed populations of pathogenic *Acinetobacter*

To test the possible transfer of antibiotic resistance in mixed populations, we first assessed if recombinants could arise in cultures of isolates harboring distinct resistance determinants. Three imipenem-resistant (Imi^R^) *A. baumannii* clinical isolates (AB5075, 40288, CNRAB1) and seven imipenem sensitive isolates either from the *baumannii* (29D2, A118, AYE, 27304, 29R1, 27024) or *nosocomialis* (M2) species were selected. Rifampicin-resistant (Rif^R^) mutants of the Imi^S^ isolates were obtained. Each of these imipenem sensitive and Rif^R^ isolates was grown for 24 hours in mixed culture with each Imi^R^ isolate. The frequency of Rif^R^/Imi^R^ recombinants in the mixed cultures was then determined. Rif^R^/Imi^R^ recombinants were detected in 19 of the 21 tested combinations with highly variable frequencies ranging from 2.10 × 10 ^-9^ to 4.82 × 10^−4^ (Fig. 1). Importantly, imipenem-sensitive isolate did not become Imi^R^ in the absence of an Imi^R^ isolate. And similarly, in the absence of Rif^R^ isolates, Imi^R^ isolates spontaneously develop resistance to rifampicin at frequencies below that of the detection limit of this assay (10^−9^). This strongly suggests that Rif^R^/Imi^R^ recombinants result from horizontal gene transfer between the mixed isolates. All tested isolates were capable of natural transformation under the tested growth conditions when presented with purified genomic DNA (gDNA) extracted from their own rifampicin resistant (Rif^R^) derivative (Fig. 1). Most isolates showed high transformability (transformation frequencies> 1 × 10^−3^) while two isolates (CNRAB1 and 27024) presented lower transformation frequencies (1 × 10^−6^). Pairing these two isolates generated only few recombinants, while pairing them with a more transformable isolate generated up to 10,000 times more recombinants, suggesting that at least one of the two isolates needs to be transformable to obtain recombinants. Yet, some combinations of highly transformable isolates were poorly productive of recombinants (40288 x AYE), suggesting that factors other than the intrinsic transformability of the isolates play a role in the production of a recombinant progeny.

**Figure 1.**
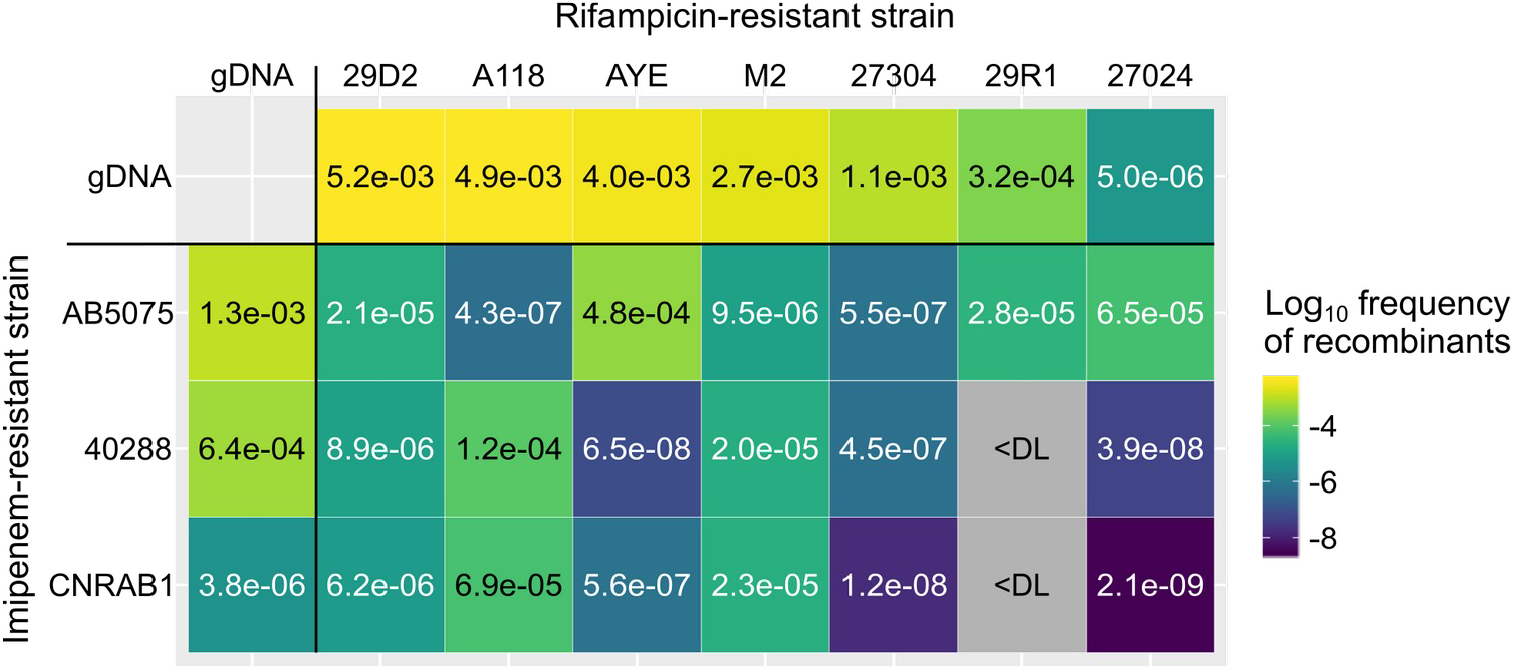
Recombinants are produced in mixed cultures of *A. baumannii* and *A. nosocomialis*. Bacterial suspensions of rifampicin-resistant (Rif^R^) isolates of carbapenem-sensitive strains (29D2, A118, AYE, M2, 27304, 29R1 and 27024) were mixed with the carbapenem-resistant (Imi^R^) clinical isolates AB5075, 40288 or CNRAB1 using cultures adjusted to an OD_600nm_ of 0.01. The mixture (2.5 µL) was deposited on the surface of tryptone-NaCl medium and incubated overnight at 37°C. Rif^R^ and Imi^R^ recombinants were determined after 24h of mixed culture. Recombinant frequencies represent the ratio of Rif^R^ and Imi^R^ colony-forming units (CFUs) over the total CFU counts. Frequency of recombinants are presented as a heat map with indicated average frequencies from two or three biological replicates with three technical replicates each. <DL, below detection limit (∼10^−9^). Top row and outermost left column show the transformation frequency displayed under the same condition by the unmixed isolates, tested by incubating the original isolates with genomic DNA (gDNA) extracted from their rifampicin (Rif^R^) derivatives. The transformation frequencies (ratio of Rif^R^ CFUs by total CFUs counts) are represented by a color gradient from blue to yellow. Data shown are the mean of natural transformation assays done in two or three biological replicates with three technical replicates each.

### Pathogenic *Acinetobacter* rapidly acquire carbapenem resistance by natural transformation in mixed culture

The conditions under which we observed the rise of recombinants are permissive to natural transformation, yet we could not rule out other mechanism of horizontal gene transfer. Natural transformation has been previously reported to occur during exponential growth in *A. baumannii* (27, 28). Therefore, we sought to determine whether the kinetics of increasing recombinant numbers under mixed culture are consistent with a role of natural transformation. Focusing on the M2 × 40288 combination of two pathogenic *Acinetobacter* strains, we could detect recombinants as early as 3 hours after mixing the two isolates (Fig. 2A). The frequency of recombinants reached a plateau at 6 hours. The kinetic of emergence of recombinant is strikingly similar when we provided the M2 isolate with genomic DNA of 40288, further indicating that recombinants emerge by natural transformation (Fig. 2A). Indeed, in a 4 hours-long mixed culture experiment, addition of DNAse I reduced the size of the recombinant population by two orders of magnitude (Fig. 2B). To further validate the role of natural transformation, we tested the emergence of recombinants by Δ*comEC::aacC4* derivatives, which are impaired for the import of DNA. The association of the M2 strain with the 40288 Δ*comEC* mutant did not alter the rate of recombinants (Fig. 2C). However, inactivating *comEC* in the M2 strain completely abolished the production of recombinants, indicating that the recombinants are formed by M2 bacteria which have acquired the carbapenem resistance by natural transformation. Similarly, the results demonstrate that the AYE and 27024 strains use natural transformation to acquire the carbapenem resistance of AB5075 (Fig. 2C). Interestingly, carbapenem-resistant transformants of 27024 are produced at rates higher than the transformation rate of 27024 when tested with its own gDNA conferring resistance to rifampicin (Fig. 1). This suggests that acquisition of carbapenem resistance is more effective than the acquisition of the mutation in the *rpoB* gene conferring resistance to rifampicin. The mutations in this housekeeping gene are probably highly costly in terms of fitness (29, 30) which may explain the absence of recombinants resulting from the acquisition of the rifampicin resistance by transformable carbapenem-resistant isolates. In conclusion, in mixed populations of *Acinetobacter*, carbapenem resistance can rapidly and efficiently spread to susceptible isolates by natural transformation.

**Figure 2.**
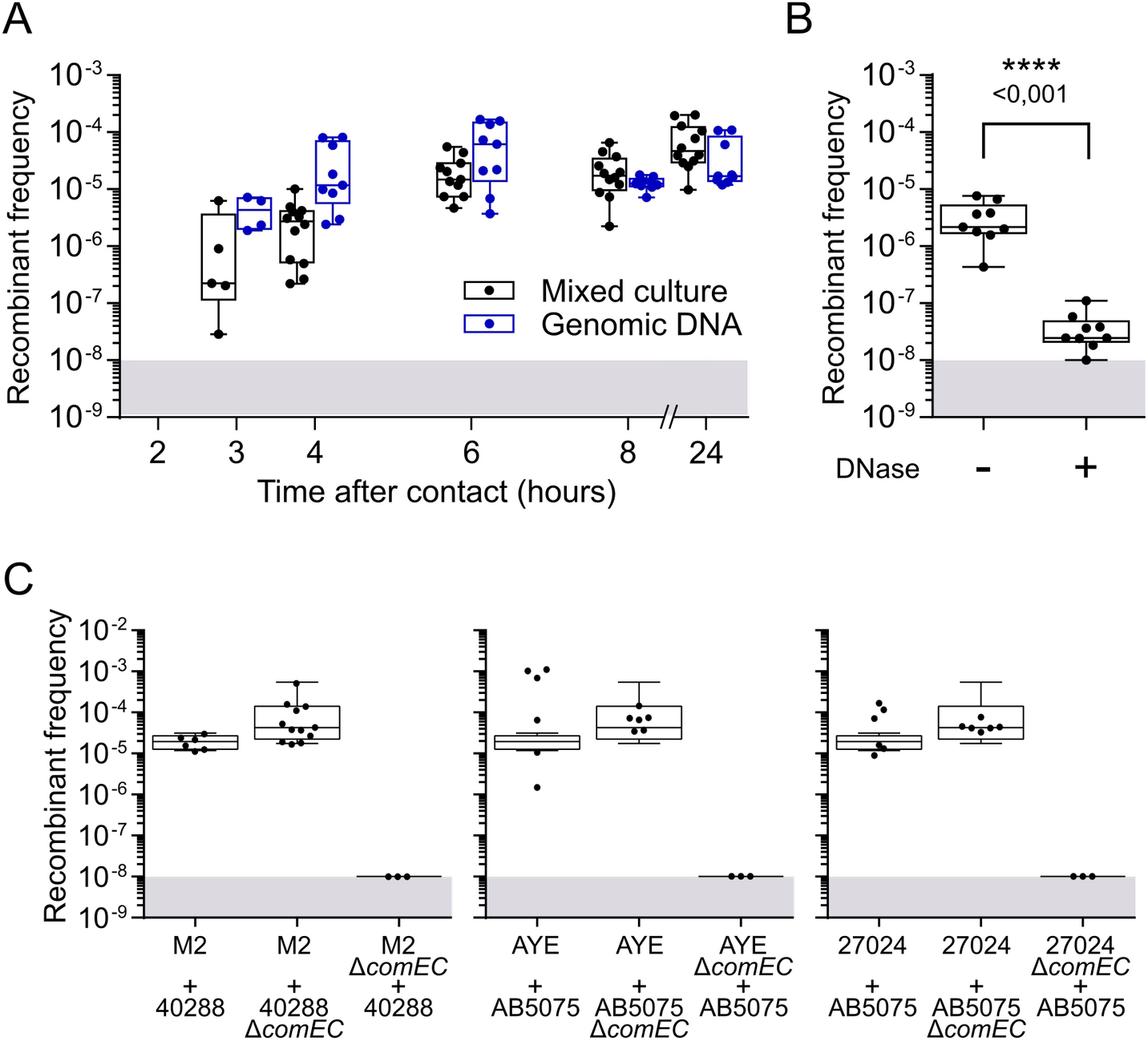
Acquisition of carbapenem resistance by natural transformation in a mixed culture. A. Kinetic of emergence of carbapenem and rifamipicin-resistant recombinants in a mixed culture of M2 Rif^R^ and 40288 (Imi^R^). Bacterial suspensions of M2 Rif^R^ at an OD_600nm_ of 0.01 were mixed with either an equal volume of a suspension of 40288 at an OD_600nm_ of 0.01 or with gDNA extracted from 40288 (at 200 ng/µL). The mixture (2.5 µL) was deposited on the surface of tryptone-NaCl medium and incubated overnight at 37°C. Recombinant frequencies represent the ratio of Rif^R^ and Imi^R^ colony-forming units (CFUs) over the total. The limit of detection (10^−8^) is indicated by the grey area. B. Sensitivity of recombinants frequencies to DNAse I treatment under the same conditions as in A, with a mixed culture of 6 hours. C. Recombinants emerge by the ComEC-dependent acquisition of carbapenem resistance by susceptible isolates. Rifampicin-resistant (Rif^R^) isolates of M2, AYE, 27024 or their *comEC* mutants were mixed with the carbapenem-resistant (Imi^R^) clinical isolates 40288, AB5075 or their *comEC* mutants. Rif^R^ and Imi^R^ recombinants were determined after 24h of mixed culture. The boxplots represent the distributions of Rif^R^ and Imi^R^ recombinants divided by the total CFU count (recombinant frequencies). A horizontal line representing the median. The limit of detection (10^−8^) is indicated by the grey area. In panel B, statistical analysis was conducted using the non-parametric Mann-Whitney-Wilcoxon test (two tailed). p-values are indicated.

### Chromosomal DNA transfer is stimulated by T6SS-killing activity, but primarily relies of contact-independent spontaneous release of DNA

The essential role of natural transformation in the acquisition of carbapenem resistance suggests that chromosomal DNA of the carbapenem-resistant isolate is released during the mixed culture. In *Vibrio cholerae*, the Type VI secretion system (T6SS) mediates the killing of non-kin cells, promoting the DNA release from prey cells which is then imported by natural transformation by the predating bacteria (31). *Acinetobacter baylyi* also produces a T6SS to kill preys, allowing the uptake of their DNA (32). In order to determine the contribution of T6SS-dependent killing on the chromosomal DNA transfer in the *Acinetobacter* mixed culture, we examined the combination of strains 40288 and M2, the latter being known for producing an active T6SS in LB medium (33). Under these conditions, we confirmed that M2 efficiently kills the 40288 strain, with a reduction of the population of 40288 by four orders of magnitude (Fig. S1A). A Δ*hcp* mutant of the M2 strain could not kill 40288, demonstrating that killing is indeed T6SS-dependent (Fig. S1A). However, in the condition in which we observed transformation-dependent chromosomal transfer, the T6SS of M2 is presumably poorly active with titers of viable 40288 cells comparable when mixed for 4 hours with either M2 or its Δ*hcp* derivative (Fig. S1B). Yet, inactivating the T6SS of M2 slightly lowered the rate of observed recombinants at 4 hours, and reduced it by one order of magnitude at 24 hours (Fig. 3A). This indicates that the residual activity of the T6SS contributes to DNA release and the acquisition of chromosomal DNA by the M2 strain. Yet, even in the absence of a functional T6SS, the rates of recombinants remained high, at a frequency of 10^−5^. We thus hypothesized that DNA is also released independently of the contact-dependent killing by the T6SS. To test this hypothesis, we collected culture supernatant of 40288 at 4 hours in the absence of M2. The M2 strain presented with this spent medium could acquire the carbapenem resistance almost as efficiently as during the mixed culture with 40288 or when presented with purified genomic DNA of 40288 (Fig. 3B). No carbapenem-resistant recombinants were observed when the spent medium was treated with DNase I or if the M2 strain is inactivated for DNA uptake (Δ*comEC*). This demonstrated that contact-independent release of DNA is sufficient to promote acquisition of chromosomal DNA by non-kin cells.

**Figure 3.**
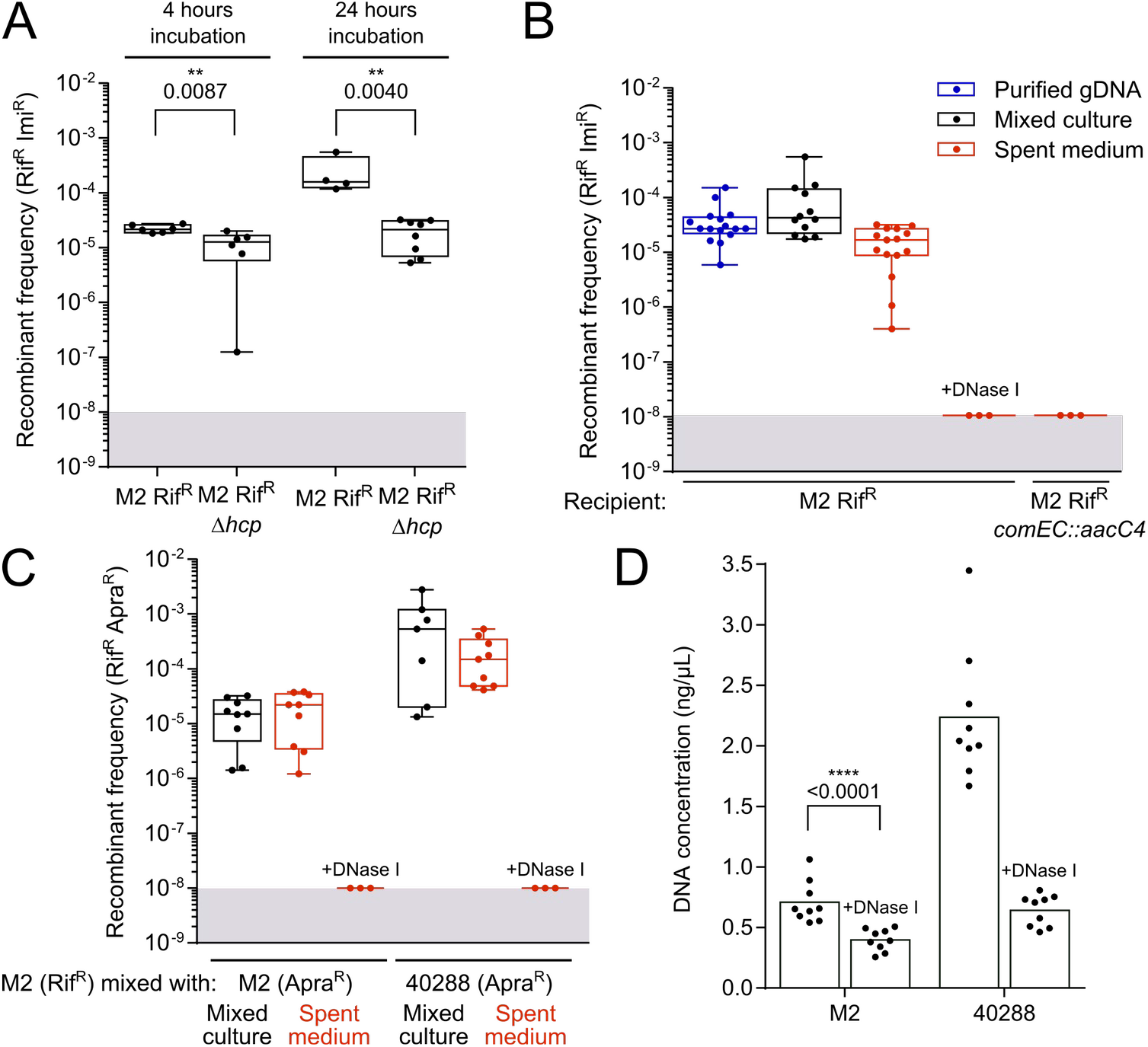
Contribution of contact-dependent T6SS-killing activity and contact-independent DNA release in the acquisition of the carbapenem resistance in mixed cultures. A. Contribution of the T6SS in the frequency of recombinants generated in a mixed culture of 40288 and M2 Rif^R^. 40288 was mixed with the M2 Rif^R^ strain or the M2 Rif^R^ Δ*hcp* mutant defective for T6SS activity. Imipenem-resistant transformants of the M2 strain were determined after 4 hours or 24 hours of mixed culture. Transformation frequencies represents the ratio of M2 Rif^R^ Imi^R^ CFUs over the total count of M2 Rif^R^ CFUs. B. Contribution of contact-independent DNA release by 40288 in the acquisition of the imipenem resistance by M2 in mixed culture. Bacterial suspensions of M2 Rif^R^ at an OD_600nm_ of 0.01 were mixed with an equal volume of either genomic DNA extracted from 40288 (at 200 ng/µL), a suspension of 40288 at an OD_600nm_ of 0.01 of with or a filtered spent medium of a 4-hours culture in tryptone-NaCl medium of 40288. The mixture (2.5 µL) was deposited on the surface of tryptone-NaCl medium and incubated for 6 hours at 37°C. Recombinant frequencies represent the ratio of Rif^R^ and Imi^R^ colony-forming units (CFUs) over the total. C. Contact-independent DNA release promotes both intra- and inter-strain recombination. The frequency of intra-strain recombinants from M2 Rif^R^ and M2 Apra^R^ strains was dermined in mixed culture or when M2 Rif^R^ is exposed to the spent medium of the M2 Apra^R^ strain. To test for inter-strain recombination, M2 Rif^R^ was mixed with 40288 Apra^R^ or with the spent medium from that strain. Recombinant frequencies represent the ratio of Rif^R^ and Apra^R^ colony-forming units (CFUs) over the total. D. Fluorescence-based quantification of DNA present in the spent medium of strains M2 and 40288. Samples treated with DNAse I represent the detection limit of such assay (∼0.4 ng). When displayed p -values of statistical analysis were obtained using the non-parametric Mann-Whitney-Wilcoxon test (two tailed). The limit of detection (10^−8^) is indicated by the grey area.

Given that DNA release is independent on the presence of another strain, DNA release and transformation may also occur between individuals of the same strain. We tested this possibility by looking at the ability of M2 Rif^R^ individuals to acquire the apramycin resistance marker (Apra^R^) of a M2 Δ*comEC* mutant. In this mixed culture setup, Rif^R^/Apra^R^ recombinants occurred at a frequency of 10^−5^ (Fig. 3C). The same frequency is observed when using spent medium of the M2 Apra^R^ Δ*comEC* mutant to transform M2 Rif^R^. Together, this shows that M2 cells spontaneously release DNA that can serve to transform other cells of the population. Yet, in a mixed culture setup, M2 acquired the apramycin resistance marker from 40288 Δ *comEC* at higher rates than from itself (M2 Δ*comEC*) (Fig. 3C). The same result is observed with the spent medium, raising the possibility that the DNA released by 40288 is more potent to produce transformants. Given that M2 can be transformed at the same efficiency with the M2 or 40288 gDNA (Fig. S2), we hypothesized that 40288 simply releases more DNA than the M2 strain. Indeed, quantification of the DNA concentration in the spent medium from a 4-hours culture of both strains, showed that 40288 releases at least 3 times more DNA than the strain M2 (Fig. 3D). Yet, and surprisingly, quantities of DNA in spent media remain very low, with concentrations of around 2 ng/µL for the 40288 strain and 0.7 ng/µL for the M2 strain, which was almost as low as the detection limit (0.4 ng/µL, obtained with DNase I-treated samples). However, this low amount collected at 4 hours of growth is sufficient to generate transformants at frequencies comparable to those obtained with about 100 times larger amounts of purified gDNA in a 6-hours window (Fig. 3B). Overall, the data indicates that the amount of spontaneously released DNA is the major determinant of the rate of recombinants generated from individuals of the same or different strains. In conclusion, the T6SS-dependent and contact-independent release of small amount of genomic DNA by pathogenic *Acinetobacter* fuels horizontal gene transfer and the acquisition of antibiotic resistance genes.

### Acquisition of resistance can result from the horizontal transfer of large genomic islands

To characterize the chromosomal transfer of the imipenem resistance at a genetic level, we sought to identify the imipenem resistance determinant in the 40288 donor strain and in transformants. 40288 is a carbapenem-resistant strain isolated from a diseased animal and carries the *bla*_OXA-23_ carbapenem-resistance resistance gene (34). An initial assembly of the 40288 genome using Illumina reads resulted in 54 contigs, and the *bla*_OXA-23_ carbapenem-resistance gene was found in a 2.4 kb contig, but whose exact location in the genome remained unresolved. A hybrid assembly combining the Illumina reads with long reads (Oxford Nanopore technology) resulted in the complete genome of 40288 showing a single circular contig of 4,084 kb. The genome revealed the presence of the *bla*_OXA-23_ gene as part of the 4.8-kb Tn*2006* transposon inserted at two distinct locations. One Tn*2006* transposon is within the large AbaR4 island of 16-kb inserted in the *comM* gene, while the other copy is inserted in the beginning of the *vgrG* gene, encoding a T6SS component. This gene, in a putative operon with a LysM domain effector, is hereafter referred as *vgrG3* (35). Insertion of the Tn*2006* into *vgrG3* gene did not impair T6SS-mediated killing, as introduction of this insertion in the *vgrG3* gene of the M2 strain did not alter its T6SS-medated killing of *E. coli* and *A. baumannii*, contrarily to a Δ*hcp* mutant (Fig. S3).

Importantly, we found that either copy of *bla*_OXA-23_ genes confers the same level of resistance to imipenem to the M2 strain (Table S3). However, the rate of their horizontal transfer may be affected by their genetic context. Acquisition of the *bla*_OXA-23_ gene from *vgrG3*::Tn*2006* requires to incorporate 4.8 kb of heterologous sequence, while acquisition of the other copy would involve the acquisition of the whole AbaR4 island of 16.6 kb. When genomic DNA from 40288 is provided to the M2 strain, the vast majority of imipenem-resistant transformants results from the acquisition of *vgrG3*::Tn*2006*, with only 5 % of transformants acquiring the 16 kb-long AbaR4 island in *comM* (Fig. 4A). In contrast, under the mixed culture condition, this percentage increased 5-fold (Fig. 4A), suggesting that the mixed culture allows the acquisition of large DNA fragments. To further determine the length of imported DNA molecules under these conditions, we characterized by whole genome sequencing the extent of DNA recombination which had occurred in the M2 transformants. To this end, fifteen imipenem-resistant transformants of M2 obtained from mixed cultures with 40288 were analyzed: seven resulting from the acquisition of *vgrG3*::Tn*2006* insertion and eight from the acquisition of *comM*::AbaR4. We used the nearly 300,000 SNPs between the two strains to identify the parts of the 40288 genome which had been incorporated in the M2 genome (Fig. S4A). The succession of acquired SNPs revealed discontinuous recombination tracts, on each sides of the acquired heterologous sequence (Fig. S4A). The discontinuous tracts most likely reflect multiple strand invasion events from the same imported DNA molecule (36). We thus used the position of the outermost acquired SNP to estimate the length of the DNA molecules of 40288 which was imported by the M2 recipients (Fig. 4B). We estimated that recombinants which have acquired Tn*2006* in *vgrG3* had imported DNA fragments ranging from 13 to 77 kb while those acquiring AbaR4 had imported DNA fragments ranging from 27 to 89 kb (Table S5). Thus, transformants resulted from the recombination of DNA molecules much larger that the acquired heterologous sequences. This suggests that the mixed culture is prone to allow the acquisition of large heterologous sequences such as those of genomic islands.

**Figure 4.**
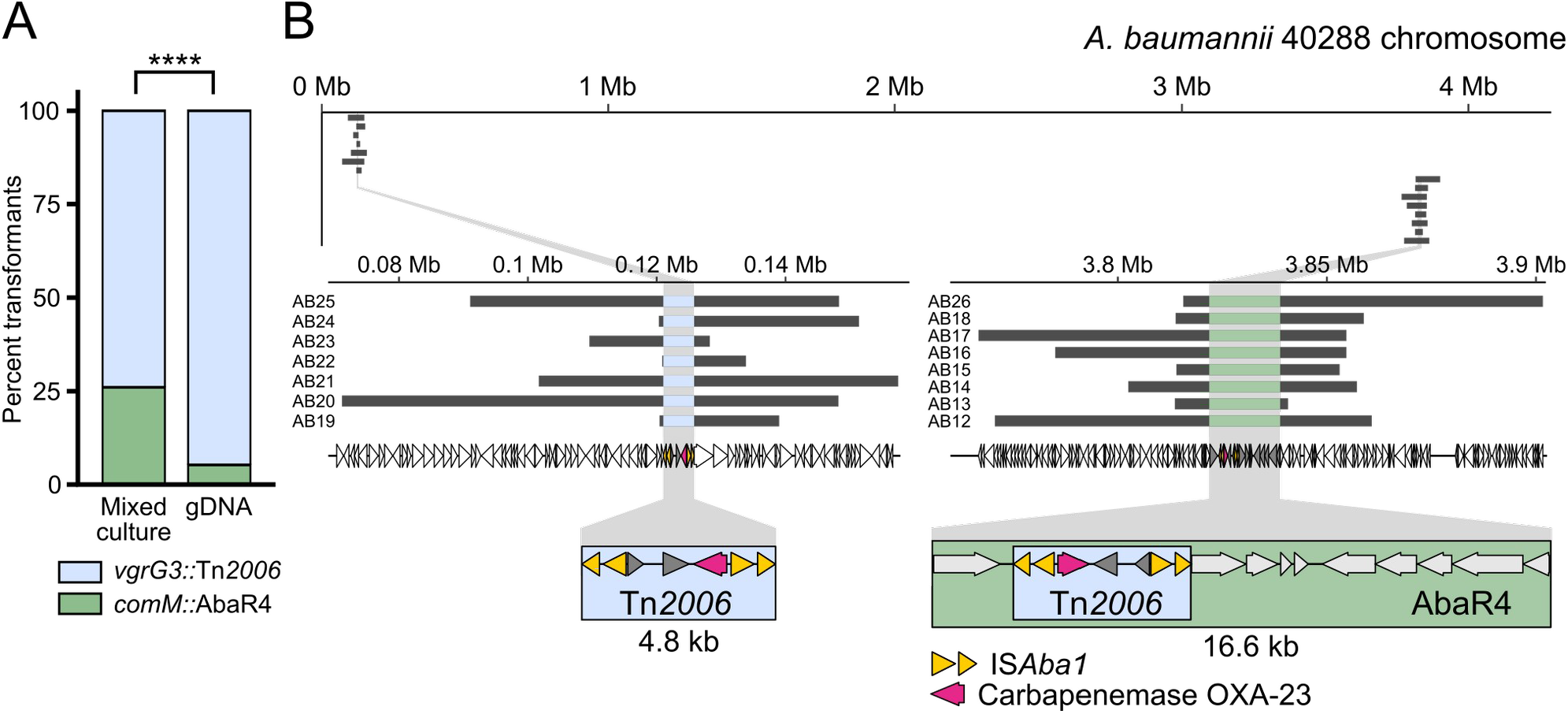
Genomic analysis of the acquisition of imipenem resistance. A. PCR-based analysis of the chromosomal location of the Tn*2006* transposon and associated *bla*_*OXA-23*_ gene conferring resistance to imipenem. One hundred Imi^R^ and *bla*_*OXA-23*_ PCR positive CFUs from 2 independent transformation assays were analyzed by PCR probing insertion in the *comM* gene. Statistical significance was calculated using Pearson’s Chi-squared test returning p-value <0.0001 (****). B. Graphical representation of the chromosome of the 40288 strain (thin black line) and location of the acquired DNA fragments (light grey thick lines) by transformants of the *A. nosocomialis* strain M2 during mixed culture with 40288. Bottom left and right panels represent close-up of the regions in which the *bla*_*OXA-23*_ gene is acquired as part of the Tn*2006* element (blue box) in the *vgrG3* gene and as part of the Tn*2006* element within the AbaR island (green box) inserted in the *comM* gene. Acquired regions were determined by sequencing the genomes of imipenem-resistant recombinants of M2 which had acquired *bla*_*OXA-23*_ gene on a Novaseq instrument. Variant calling and identification of converted markers (SNPs of 40288 acquired by M2) were used to delineate the acquired regions (see Material and Methods).

The AbaR4 island is one of the many representatives of resistance islands that can be found in *A. baumannii* clinical isolates (9). The largest resistance island described so far, the 86 kb-long AbaR1, is found in the AYE strain (8). We thus tested whether this island could be acquired by the M2 strain. In a previous work, we showed that the AbaR1 island conferred resistance to aminoglycosides, tetracycline and to some beta-lactams (37). We thus used the tetracycline resistance as a marker for the transfer of AbaR1 from the *A. baumannii* strain AYE (donor strain) to the *A. nosocomialis* strain M2 (recipient strain). To test the acquisition of the AbaR1 by M2, we used the rifampicin resistant derivative of strain M2 and a derivative of the strain AYE impaired for natural transformation (Δ*comEC::aacC4*). The mixed culture condition resulted in recombinants emerging at a frequency of about 10^−7^ (Fig. 5A). Noteworthy, only few transformants were detected when using purified genomic DNA, confirming that the mixed culture brings together conditions favorable to the transfer of large resistance island (Fig. 5A). The resulting transformants displayed the resistance profile expected from the known resistance associated with AbaR1 (Table S4), indicating that the entire 86 kb-long island had indeed been acquired. Genome sequencing of three tetracycline resistant transformants confirmed that the entire AbaR1 island (86 kb) was inserted into the genome of the recipient cells (Fig. 5B and Fig. S4B). The recombination tracts flanking the *comM* locus indicated that the imported DNA molecules ranged from 112 to 123 kb (Fig. 5B and Fig. S4B). Taken together, these results demonstrate that recombinants can emerge from the encounter of *Acinetobacter* isolates, even if their genome display relatively low sequence identity (90%). The interbacterial transfer can result in the efficient recombination of homologous regions and the acquisition of large heterologous segments up to 86 kb, such as AbaR islands conferring resistance to multiple antibiotics, including to carbapenems.

**Figure 5.**
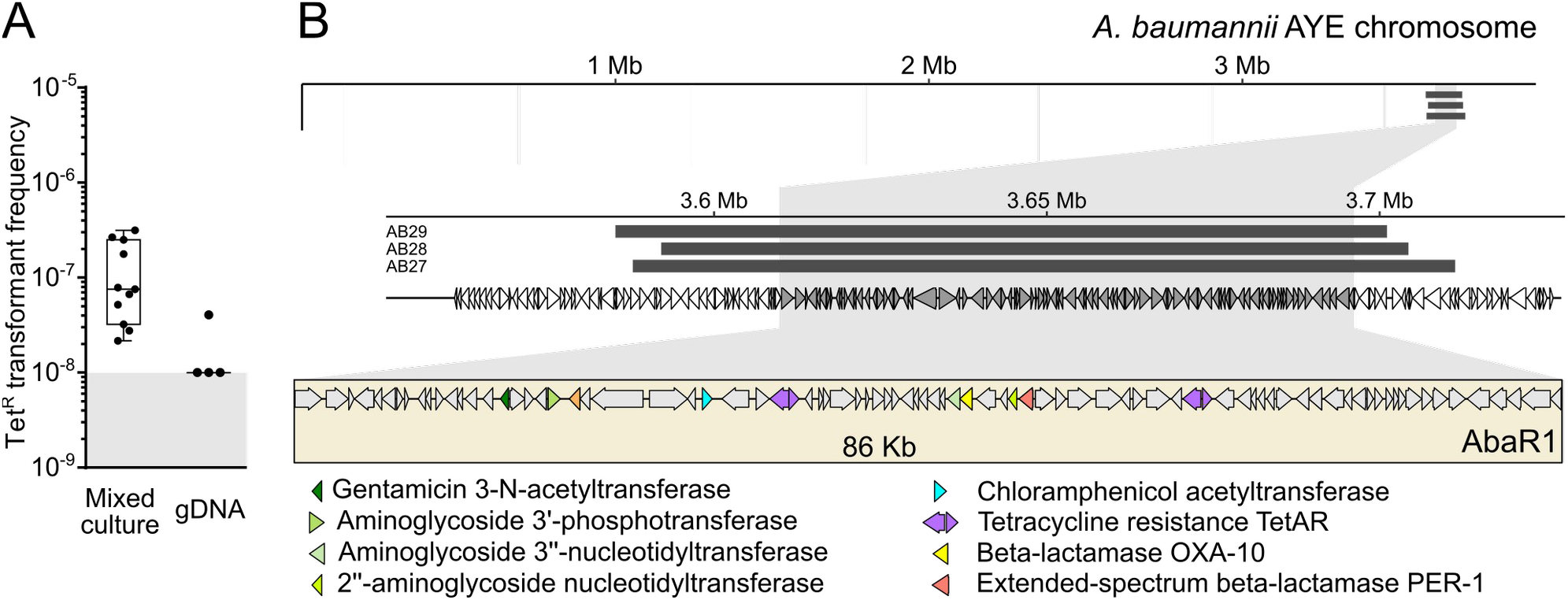
Genomic analysis of the acquisition of the AbaR1 island. A. Transformation frequencies of the acquisition of tetracycline resistance carried by the AbaR1 island of the AYE strain, in a mixed culture setup between the AYE strain impaired for natural transformation (*comEC::aacC4*) and the M2 Rif^R^ strain and by supplying M2 Rif^R^ with genomic DNA extracted from AYE. B. Graphical representation of the chromosome of the AYE strain (thin black line) and location of the acquired DNA fragments (light grey thick lines) by transformants of the *A. nosocomialis* strain M2 during mixed culture with AYE. Bottom panel, close-up view of the acquired fragments and the AbaR1 island. The genomes of tetracycline-resistant recombinants of M2 were sequenced. Variant calling and identification of converted markers (SNPs of AYE acquired by M2) were used to delineate the acquired regions (see Material and Methods).

## Discussion

To gain insight into the mechanism leading to emergence of carbapenem and more broadly multidrug resistance in pathogenic Acinetobacter sp. we devised an experimental system to monitor emergence of recombinant progeny and diffusion of antibiotic resistance between multiple combinations of pathogenic *Acinetobacter* strains. We observed that resistance is efficiently acquired within mixed populations of *A. baumannii* strains and even between *A. baumannii* and *A. nosocomialis*. With the panel of strains tested, we found an overall apparent correlation between the level of transformability of the isolates and the frequency of emergence of recombinant progeny. Investigating in more details a specific mixed population, we demonstrated that rapid gene transfer occurs through natural transformation in communities of sessile cells leading to antibiotic resistance. Resistance genes were either carried by composite transposon (Tn*2006* carrying *bla*_OXA-23_) but also by large resistance island (AbaR4 and AbaR1). So far, demonstrations of gene acquisition by natural transformation in *A. baumannii* and *A. nosocomialis* were obtained using purified DNA (23–28). Our results indicate that this later form of DNA is a poor substrate for transformation in comparison to the DNA released in the extracellular milieu during growth or upon bacterial interaction. Mixed culture allowed a transfer of up to 123 kb from strain *A. baumannii* 40288 to *A. nosocomialis* corresponding to over 3% of the recipient strain’s genome. Our results corroborate seminal observations in *Streptococcus pneumoniae* in which mixed sessile populations were found to support larger recombination events (up to 29 kb) (38). Similarly, interaction of *V. cholerae* strains also promotes transfer of large genomic region (up to 168 kb) (39). Importantly, our experiment recapitulate the observed recombination events associated with the acquisition of AbaGRI3 by a strain from the global clone 1 (GC1) originated from the GC2 (14). Acquisition by natural transformation would therefore explain the acquisition of AbaRs described in GC2 strains (40) but also in other *Acinetobacter* species from the *A. baumannii*-*A. calcoaceticus* complex, namely, *A. nosocomialis, haemolyticus, pitii*, and *seifertii* (9, 41, 42). Our experimental observation leads to reconsider the role of natural transformation in the acquisition of large islands of resistance by *A. baumannii* previously attributed to generalized transduction based on presumed limited size of acquisition enabled by natural transformation (20). Transfer of very large resistance island such as AbaR1 are less frequent as the odds of importing an intact DNA are likely lower for long DNA fragments. Moreover, acquisition of new genetic material resistance genes has a potential cost on bacterial replication (43). In the specific case of AbaR1, importing numerous antibiotic and heavy metal resistance genes may be detrimental for the recipient strain with a temporary fitness cost that would affect the apparent transformation frequency (44, 45). If natural transformation is indeed a route of diffusion of large resistance island, these non-exclusive hypothesis would explain why resistance islands are mainly of about 20 kb-long and carry a limited number of antibiotic resistance genes (9).

We investigated the role of T6SS in the transfer of resistance upon bacterial interaction. Indeed, this competition mechanism was an evident culprit to trigger the release of DNA from prey cells, as described for *A. baylyi* or *V. cholerae* (31, 32, 39). Although we found that T6SS is partially involved in acquisition of resistance genes, we found however that T6SS mediated killing is potentially impaired in conditions compatible with natural transformation. Our results corroborate previous findings that T6SS development is dependent on growth conditions with the observation of a reduced T6SS activity determined by Hcp secretion in minimal medium for *A. nosocomialis* M2 strain (46). Contribution of T6SS in DNA release and diffusion of resistance determinants may however depend of strains combination, as T6SS activity was shown to be highly dependent of the strains in competitions (35, 47, 48). In addition, our results show that the direct negative interaction between cells is not the sole mechanism promoting the release of DNA and the subsequent the acquisition of the resistance determinant. We found that the contact-independent release of DNA during growth is sufficient to mediate high rates of acquisition of resistance through natural transformation. Our results indicate that pathogenic *Acinetobacter* strains may release varying amount of DNA in the extracellular medium. In addition to passive release by autolysis upon cell death, several mechanisms of active DNA release in the extracellular compartment had been described in other transformable species. This release could be consecutive to a concerted lysis of a specific subpopulation. Indeed, competent cells of *S. pneumoniae* are able to recognize and kill non-competent kin cells leading to the lysis of up to 30% of the population (49, 50). In *Neisseria gonorrhoeae*, genomic DNA is actively secreted through a Type 4 Secretion System by the donor strain delivering a more potent substrate of transformation than DNA released upon autolysis (51, 52). In *Pseudomonas aeruginosa*, DNA release is dependent on a cryptic prophage endolysin that is required from biofilm formation where transformation can also occur (53– 55). We found that DNA concentration in the culture supernatant of *A. baumannii* strain 40288 was about 2 ng/µL and a potent substrate for transformation, a result consistent with concentration measured in *P. aeruginosa* strain PAO1 culture (56). Extracellular DNA is also a critical component of biofilm matrix (57) and treatment with DNAse leads to severe biofilm alteration for several pathogenic bacteria, including *A. baumannii* (58). Inhibition of DNA release could therefore mitigate antibiotic resistance transmission by natural transformation and biofilm formation of this nosocomial pathogen. However, results obtained in *P. aeruginosa* demonstrates that biofilm-associated DNA is highly structured by nucleoproteins in a lattice that may partially impair its availability for transformation (59, 60). It is conceivable that level of DNA release may variate between strains of *A. baumannii* and thereby influence the diffusion of resistance gene and their acquisition by neighboring cells.

Altogether, this work highlights the importance of one of the major HGT mechanisms in bacteria, natural transformation, in the acquisition by *A. baumannii* of resistance to clinically relevant antibiotics. By naturally releasing high quality DNA in their environment, some strains of *A. baumannii* may be highly potent donors for the uptake and recombination of large DNA fragments by natural transformation. This mechanism may enable a rapid and direct acquisition of multiple antibiotic resistance genes by *A. baumannii* and explain the evolution by recombination observed at the genomic scale.

## Materials and Methods

### Bacterial strains and growth conditions

The bacterial strains or strains used in this study are listed in supplementary table S1. Unless specified, *Acinetobacter baumannii* isolates and strains were grown in lysogeny broth (LB) (Lennox). All experiments were performed at 37°C. Antibiotic concentrations were: 15 µg/mL for tetracycline, 1.6 µg/mL for imipenem (always spread extemporaneously on LB agar plate) and 100 µg/mL for rifampicin.

### Construction of bacterial strains

All the oligonucleotides used in this study for genetic modification are listed in supplementary table S2. Gene disruptions were performed using a scarless genome editing strategy described previously (37). Overlap extension PCR were used to synthesize large chimeric DNA fragments carrying the selection marker flanked by 2 kb fragments that are homologous to the insertion site. The oligonucleotides used for strain construction are listed in Table S2. The PCRs were performed with a high-fidelity DNA polymerase (PrimeStarMax, Takara).

### Detection of recombinants in mixed cultures

Isolates to be tested for the production of recombinants were grown overnight on LB agar plates, then inoculated and grown in 2 mL of LB until the cultures reach an OD _600nm_ of 1. The bacterial broths were then diluted to an OD_600nm_ of 0.01 in PBS. Then equal volumens of bacterial suspensions were mixed by pairs, each consisting of an imipenem-resistant (Imi^R^) and rifampicin-resistant (Rif^R^) isolate. The mixture (2.5 µL) was deposited on the surface of 1 mL of tryptone-NaCl medium solidified with 2% agarose D3 (Euromedex) poured in 2 mL micro tubes or in wells of 24-well plates and incubated overnight at 37°C. Depending on the method used for plating, 200 or 400 μl of DPBS, were added to the tubes which were vigorously vortexed to resuspend the bacteria. In 24 well plates, 200 or 400 μl of DPBS and glass beads were added to the wells, the plates were then shaken to resuspend the bacteria. Suspension were plated on LB agar plates without antibiotics and LB agar plates containing rifampicin (100 µg/mL) and imipenem (1.6 µg/mL) either with beads or easySpiral Pro (Interscience). Recombinants frequencies were determined through calculation of the ratio of the number of CFUs on rifampicin and imipenem plates, to the total number of CFUs on plates without antibiotics.

### Transformation assay

After overnight incubation at 37°C on LB agar, the strains were grown in 2 mL of LB until OD_600nm_ of 1. The bacterial broths were then diluted to an OD_600nm_ of 0.01 in DPBS. Then bacterial suspensions of the recipient strain were mixed with an equal volume of either genomic DNA extracted from the donor strain (concentration 200 ng/µL), or a bacterial suspension of the donor strain at an OD_600nm_ of 0.01 for mixed culture, or with a filtered spent medium of a 4-hours culture in tryptone-NaCl medium (5 g/liter of tryptone, 2.5 g/liter NaCl) of the donor strain. The mixture (2.5 µL) was deposited on the surface of 1 mL of tryptone-NaCl medium solidified with 2% agarose D3 (Euromedex) poured in 2 mL micro tubes and incubated overnight at 37°C. The next day, bacteria were recovered by resuspension in 300 µL of PBS, serial diluted and spread on LB agar plate without antibiotic or supplemented with the appropriate antibiotic. In experiments including DNAse I treatment, 0.6U of DNAse I (Sigma) were added per microliter of the mixed culture. All the transformation assays were performed on at least two separate occasions. On each occasion, three independent transformation reactions were conducted (three different bacterial cultures). All the independent data points are plotted. As normality of the distribution of transformation frequency does not apply for transformation frequency analysis, non-parametric tests were performed (Mann-Whitney-Wilcoxon).

For kinetics of transformation, bacteria were recovered from the solidified tryptone-NaCl medium at various times points: 0 and 30 min after contact, then one point every hour from 1 to 8 hours after contact and one final point at 24 hours after contact. For each point, bacteria were harvested from an independent sample and plated as previously described.

### Determination of antagonistic activity between strains

T6SS-dependent killing activity of the M2 strain were determined in LB medium and in conditions of mixed culture, as previously described (33). Strains were grown 2 h at 37°C in the indicated liquid medium, then were diluted to an OD_600nm_ of 0.4. The bacterial suspensions were then mixed at a ratio of 10:1 (predator:prey) and 10 µL of the mixed suspension were spotted on a dried plate and incubated at 37°C during 4h. Bacteria were then collected from the plate, resuspended and serial dilution were plated on selective plate to determine the titer of prey bacteria. Plates containing kanamycin and imipenem were used to selectively recover the prey bacteria *E. coli* JW2912 and *A. baumannii* 40288, respectively.

### DNA measurement in spent media

Bacteria were prepared as for transformation assay lasting 4 h at 37°C in 2 mL-eppendorf tubes. After the 4h incubation, the cells were harvested with 300 µL of sterile water. As control, 25 µL of the bacterial suspension were used for serial dilutions in 96-well plates for numbering on LB agar with or without supplements to verify the survival and M2 transformation efficiencies. DNA quantification in the spent medium of the donor was performed through fluorescent labelling with Quant-iT reagent (Invitrogen Qubit dsDNA HS Assay Kit, Thermo Fisher Scientific). Fluorescence was measured with Infinite M200 PRO (TECAN) and Magellan 7.1 SP1 (TECAN) and DNA concentration was calculated by comparison with Quant-iT standards (Invitrogen Qubit dsDNA HS Assay Kit, Thermo Fisher Scientific) in liquid Tryptone NaCl medium.

### Genome sequencing, assembly, and annotations

The genome of 40288 was sequenced using the Illumina technology (paired-end 100, HiSeq instrument) and Oxford nanopore technology (rapid barcoding kit, MinIon instrument) with a read coverage of 100X. Short (Illumina) and long (Oxford nanopore) reads were used to assemble the genome with Unicycler (61). This generated a complete genome consisting of a circular chromosome of 4,084,922 bp and circular plasmid of 145,711 bp. Genes were annotated with Prokka (62), using the *A. baumannii* AB5075-UW genome as reference (63). The genome of the human clinical isolate CNRAB1 was sequenced using similar procedures. Genome sequences are available within the bioproject PRJNA741866.

### Variant-calling and detection of recombination events

The genomes of imipenem-resistant recombinants of M2 which had acquired *bla*_OXA-23_ were sequenced on a Novaseq instrument (Illumina, 2 × 150 bp paired-end, 300 bp insertion length). Sequencing reads were subsampled to match an estimated 50x coverage of donor genome using *seqtk* version 1.3-r106 (https://github.com/lh3/seqtk). Reads of recombinants were mapped to their corresponding donor genome (M2, 40288 or AYE) using *bwa mem* version 0.7.17-r1888 (64) and sorted using *samtools* version 1.10.3 (65). Donor (40288 or AYE) and recipient (M2) genome short reads were simulated using *wgsim* version 1.10 (https://github.com/lh3/wgsim) and mapped to the donor genome (40288 or AYE) using the same parameters as for transformant sequencing reads. Reads were then tallied on donor genome using *bcftools* version 1.10, to keep only sites with read coverage above 25x, where haplotype differs between donor and recipient, and match donor haplotype. These variant calling and identification of converted markers were performed using *twgt* version 0.1.0.5 (https://gitlab.com/bacterial-gbgc/twgt/-/tree/master) The resulting VCF file was then parsed using *VariantAnnotation* (66) from the Bioconductor framework version 3.10, using R version 3.6.6. Tn*2006*, AbaR4 and AbaR1 being absent from recipient genome, they cannot be detected from single nucleotide polymorphisms and short insertions/deletions. Their insertions in the transformant genome was thus confirmed via read screening: their sequence together with 1 kb of flanking sequence was sketched using *mash sketch* with default parameters; the presence of this sketches in sequencing reads was confirmed using *mash screen* version 2.2.2 (67). Recombinants carrying AbaR4 correspond to samples AB12 to AB18 and AB26. Recombinants carrying Tn*2006* in *vgrG* correspond to samples AB19 to AB25. Recombinants carrying AbaR1 are samples AB27 to 29. Raw sequencing reads are available as Sequence Read Archive (SRA) within PRJNA741866.

## Acknowledgments

We warmly thank Agnese Lupo and Marisa Haenni (Unité Antibiorésistance et Virulence Bactériennes, ANSES, Lyon, France) for sharing the clinical isolate 40288, sequencing data and antimicrobial susceptibility testing data. This work was supported by the LABEX ECOFECT (ANR-11-LABX-0048) of Université de Lyon, within the program “Investissements d’Avenir” (ANR-11-IDEX-0007) operated by the French National Research Agency (ANR). ASG and MHL were also supported by Programme Jeune Chercheur from VetAgro Sup. AL and MHL were supported by the French Agency for Food, Environmental and Occupational Health & Safety (ANSES).

