## Supplemental Material for "Interbacterial transfer of carbapenem resistance and large antibiotic resistance islands by natural transformation in pathogenic *Acinetobacter*"

**Table S1**: Bacterial strain used in this study

**Table S2**: Primers used in this study

**Table S3**: Antimicrobial susceptibility profiles of imipenem resistant transformants.

**Table S4**: Antimicrobial susceptibility profiles of tetracycline resistant transformants.

**Table S5**: Estimated length of the imported DNA molecules leading to the detected recombination tracts.

**Figure S1**: T6SS-mediated killing assay of *A. nosocomialis* strain M2 in LB medium.

**Figure S2**: T6SS-mediated killing assay of a vgrG3 impaired mutant of *A. nosocomialis* strain M2 in LB medium.

**Figure S3**: T6SS-mediated killing assay of a VgrG3 impaired mutant of *A. nosocomialis* strain M2 in LB medium.

**Figure S4**: Genomic analysis of the acquisition of imipenem resistance.

**Figure S5**: Genomic analysis of the acquisition of the AbaR1 island

**Table S1: bacterial strains and plasmids used in this study.**

| Strain | Description/phenotype | Reference |
|---|---|---|
| Recipient strains for natural transformation | | |
| M2 | *A. nosocomialis* strain M2 wild-type | (Niu et al., 2008) |
| M2 $Rif^R$ | Spontaneous *rpoB* mutant; rifampicin resistant | This study |
| M2 $Rif^R$ Δ*comEC::aacC4* | Insertion of an apramycin resistance cassette in the *comEC* gene of the M2 $Rif^R$ mutant; Non transformable and rifampicin resistant | This study |
| AYE | *A. baumannii* strain AYE wild-type | (Fournier et al., 2006) |
| AYE $Rif^R$ | Spontaneous *rpoB* mutant; rifampicin resistant | This study |
| 29D2 | *A. baumannii* nonclinical isolate 29D2 wild-type | (Wilharm et al., 2017) |
| 29D2 $Rif^R$ | Spontaneous *rpoB* mutant; rifampicin resistant | This study |
| A118 | *A. baumannii* strain A118 wild-type | (Ramirez et al., 2010) |
| A118 $Rif^R$ | Spontaneous *rpoB* mutant; rifampicin resistant | This study |
| 27304 | *A. baumannii* veterinary clinical isolate 27304 | Resapath, ANSES |
| 27304 $Rif^R$ | Spontaneous *rpoB* mutant; rifampicin resistant | This study |
| 27024 | *A. baumannii* veterinary clinical isolate 27024 | Resapath, ANSES |
| 27024 $Rif^R$ | Spontaneous *rpoB* mutant; rifampicin resistant | This study |
| Donor strains for natural transformation | | |
| 40288 | *A. baumannii* veterinary clinical isolate 40288; carbapenem resistant | (Lupo et al., |

| | | |
|---|---|---|
| | | 2016) |
| 40288 Δ*comEC::aacC4* | Insertion of an apramycin resistance cassette in the *comEC* gene; Not naturally transformable | This study |
| AB5075 | *A. baumannii* strain AB5075 wild-type; carbapenem resistant | (Jacobs et al., 2014) |
| 37986 | *A. baumannii* veterinary clinical isolate 37986; carbapenem resistant | (Lupo et al., 2016) |
| CNRAB1 | *A. baumannii* human clinical isolate CNRAB1; carbapenem resistant | CNR* |
| M2 Δ*comEC::aacC4* | Insertion of an apramycin resistance cassette in the *comEC* gene; Not naturally transformable | This study |
| AYE Δ*comEC::aacC4* | Insertion of an apramycin resistance cassette in the *comEC* gene; Not naturally transformable | This study |
| AB5075 Δ*comEC::aacC4* | Insertion of an apramycin resistance cassette in the *comEC* gene; Not naturally transformable | This study |

*Centre National de Référence de l'antibiorésistance de Besançon, France.

**Table S2: primers used in this study**

| Genetic construct | Name | Sequence 5' to 3' | Template; Annealing site |
|---|---|---|---|
| Chromosomal modifications using overlap extension PCR | | | |
| *comEC::aacC4* | comEC_UP_F | TGGTGTCGCTGTACTAATTACGGT | *Acinetobacter* genomic DNA; 2kb upstream of the *comEC* gene |
| | comEC_apra_R | GTGCCGTGCTAGTAGCACGCCCTCCCGTTCAGAGCCTCGCCCACTTTTTC | *Acinetobacter* genomic DNA; Chimeric, middle of the *comEC* gene |
| | comEC_apra_F | GGCGAGCGGTCAGCTAACCGACTCGAGTACTGGATAGGCTCTGGTTGAGTA | *Acinetobacter* genomic DNA; Chimeric, middle of the *comEC* gene |
| | Apr_Fw | CAGGCTGGGTGCCAAGCTCT | pMHL-2 (Godeux et al., 2018) ; downstream *aacC4* gene |
| | Apr_Rev | TCATGAGCTCAGCCAATCGACTGG | pMHL-2 (Godeux et al., 2018) ; upstream *aacC4* gene |
| | comEC_DW_R | CTTCAGTTCACGCATCAAGCTTGT | *Acinetobacter* genomic DNA; 2kb upstream of the *comEC* gene |
| Primers for control of the transformants using colony PCR | | | |
| | comM-For | CCACAATGGAACAAGAAGATGTCT | M2 genomic DNA; 5' of the *comM* gene |
| | comM-Rev | TTAAGAGTGATTACCTCGATAAGA | M2 genomic DNA; 3' of the *comM* gene |
| | OXA23-For | GATCGGATTGGAGAACCAGA | $bla_{OXA-23}$ gene |
| | OXA23-Rev | ATTTCTGACCGCATTTCCAT | $bla_{OXA-23}$ gene |
| | mlo-80 | CGGATCTTCGATGCTGGC | AbaR1 from AYE strain |
| | mlo-84 | GCAACGATGTTACGCAGC | AbaR1 from AYE strain |
| | 5'-J-Rev | ATGGAATGTAGTACTCTGACG | AbaR1 from AYE; AbaR junction |
| | 3'-J-For | GATTCACATCATATTCATTCGCCC | AbaR1 from AYE; AbaR junction |

| Antibiotic | Minimum inhibitory concentration (E-test)<br>40288 WT | M2 WT | M2 Rif$^R$ *comM::*AbaR4 | M2 Rif$^R$ *vgrG3::*Tn*2006* |
|---|---|---|---|---|
| IMP | >32 | 0.25 | >32 | >32 |

**Table S3: Antimicrobial susceptibility profiles of imipenem resistant transformants.** Two different transformants that acquired a distinct genetic context of the Tn*2006*-*bla*OXA-23 were tested. In M2 *comM::*AbaR4, the transposon is localized in the AbaR4 inserted in the *comM* gene. In M2 *vgrG3::*Tn*2006*, the transposon is inserted in the *vgrG3* gene encoding a T6SS element. Susceptibility of the strains to imipenem were compared by E-test strips (BioMérieux, France). The strain *Pseudomonas aeruginosa* CIP 7110 was used as control.

Inhibition zone diameter in mm

| Antibiotic | AYE WT | M2 WT | M2 Rif$^R$ | M2 Rif$^R$ *comM::*AbaR1 Clone 1 | M2 Rif$^R$ *comM::*AbaR1 Clone 2 |
|---|---|---|---|---|---|
| AMK | 8 | 29 | 28 | 15 | 14 |
| GEN | 6 | 29 | 28 | 12 | 12 |
| TOB | 6 | 26 | 25 | 9 | 7 |
| TIG | 19 | 25 | 27 | 29 | 27 |
| FOS | 8 | 6 | 6 | 6 | 6 |
| COL | 19 | 19 | 20 | 20 | 20 |
| CIP | 6 | 26 | 29 | 29 | 29 |
| PIP | 6 | 22 | 22 | 12 | 14 |
| TIC | 6 | 30 | 25 | 6 | 6 |
| IMP | 28 | 40 | 40 | 40 | 40 |
| TZP | 14 | 29 | 25 | 25 | 25 |
| TIM | 15 | 29 | 28 | 23 | 25 |
| MEM | 20 | 33 | 27 | 23 | 24 |
| CAZ | 6 | 23 | 22 | 6 | 6 |
| FEP | 6 | 26 | 25 | 6 | 6 |
| ATM | 6 | 16 | 10 | 6 | 6 |
| RIF | 13 | 17 | 6 | 6 | 6 |

Minimum inhibitory concentration (E-test)

| Antibiotic | M2 | AYE WT | AYE ΔAbaR | M2 Rif$^R$ *comM::*AbaR1 Clone 1 | M2 Rif$^R$ *comM::*AbaR1 Clone 2 |
|---|---|---|---|---|---|
| TET | 4 | 128 | 8 | 48 | 48 |

**Table S4: Antimicrobial susceptibility profiles of tetracycline resistant transformants.** Susceptibility of the strains to a panel of 17 antibiotics: amikacin (AMK), gentamicin (GEN), tobramycin (TOB), tigecycline (TIG), Fosfomycin (FOS), colistin (COL), ciprofloxacin (CIP), piperacillin (PIP), piperacillin-tazobactam (TZP), ticarcillin (TIC), ticarcillin-clavulanic acid (TIM), ceftazidime (CAZ), cefepime (FEP), meropenem (MEM), aztreonam (ATM), rifampin (RIF) and imipenem (IMP) was evaluated by disc diffusion on Muller-Hinton agar (Biorad, France) following the CA-SFM 2013 recommendations. Inhibition values were interpreted according to CA-SFM 2013 breakpoints for all antibiotics but aztreonam, for which *Pseudomonas* spp. breakpoints were used. Resistance of wild type AYE, AbaR-cured AYE mutant (AYE ΔAbaR from Gdeux et al. 2020) and M2 *comM::*AbaR1 transformants were also compared by E-test strips (bioMérieux, France) for tetracycline (TET). The strain *Pseudomonas aeruginosa* CIP 7110 was used as control for all the antibiotic susceptibility testing.

| Sample | Donor strain | Loci | First acquired SNP | Last acquired SNP | Length of imported DNA (bp) |
|---|---|---|---|---|---|
| AB25 | 40288 | Tn2006 | 91048 | 148295 | 57247 |
| AB24 | 40288 | Tn2006 | 120337 | 151404 | 31067 |
| AB23 | 40288 | Tn2006 | 109565 | 128246 | 18681 |
| AB22 | 40288 | Tn2006 | 120849 | 133850 | 13001 |
| AB21 | 40288 | Tn2006 | 101695 | 157494 | 55799 |
| AB20 | 40288 | Tn2006 | 71139 | 148223 | 77084 |
| AB19 | 40288 | Tn2006 | 120442 | 139003 | 18561 |
| AB26 | 40288 | AbaR4 | 3815631 | 3901652 | 86021 |
| AB18 | 40288 | AbaR4 | 3813832 | 3858827 | 44995 |
| AB17 | 40288 | AbaR4 | 3766851 | 3854677 | 87826 |
| AB16 | 40288 | AbaR4 | 3785214 | 3854677 | 69463 |
| AB15 | 40288 | AbaR4 | 3814000 | 3853045 | 39045 |
| AB14 | 40288 | AbaR4 | 3802492 | 3857180 | 54688 |
| AB13 | 40288 | AbaR4 | 3813642 | 3840698 | 27056 |
| AB12 | 40288 | AbaR4 | 3770779 | 3860715 | 89936 |
| AB29 | AYE | AbaR1 | 3585163 | 3701113 | 115950 |
| AB28 | AYE | AbaR1 | 3592041 | 3704339 | 112298 |
| AB27 | AYE | AbaR1 | 3587780 | 3711356 | 123576 |

**Table S5. Estimated length of the imported DNA molecules leading to the detected recombination tracts.**

SNPs acquired by the M2 recipient strain were mapped onto the donor genome and position of the first and last acquired provide an estimate of the length of the imported DNA molecule that resulted in the observed recombination tracts.

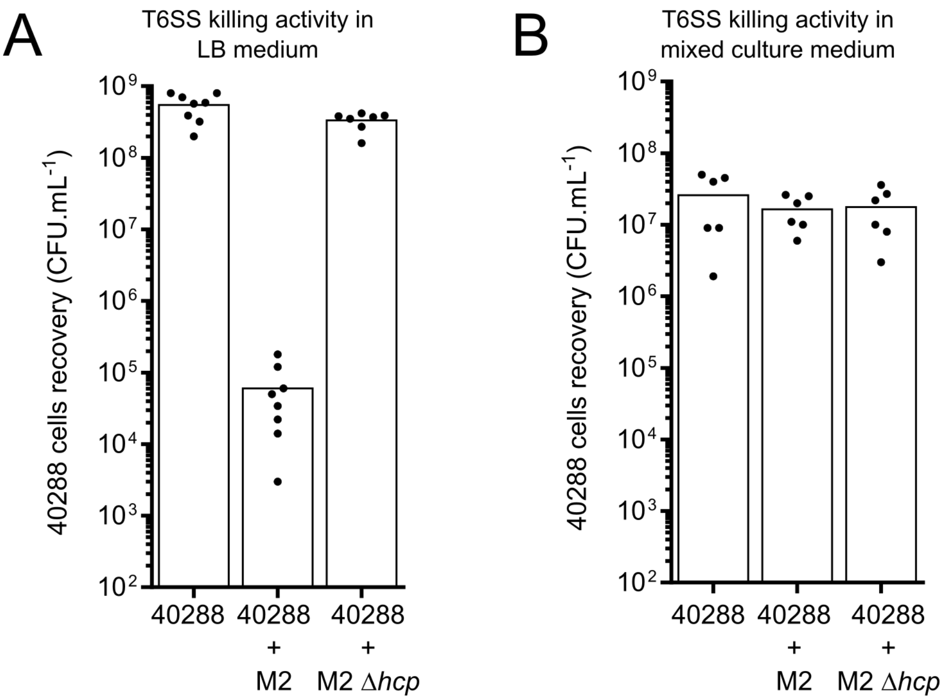


**Figure S1. T6SS-dependent killing activity of the M2 strain on the 40288 isolate in LB medium and in conditions of mixed culture.**

Recovery of the *A. baumannii* strain 40288 incubated either with the wild type M2 strain (WT) or its *hcp* gene mutant derivative (Δ*hcp*). Strains were grown 2h at 37°C in the indicated liquid medium, when ODs were below 2, they were diluted to an OD of 0.4. The bacterial suspensions were then mixed for a ratio of 10:1 (predator:prey) and 10µL of the mixed suspension spotted on a dried plate then incubated at 37°C during 4h.

A. Co-cultivation of 40288 with the M2 strain during in LB medium resulted in a 4-log decrease of the 40288 population, while inactivating the T6SS of M2 (Δ*hcp*) abolished the killing activity.

B. In the mixed culture setup used to observe the rise of recombinants, mixed culture of 40288 with M2 during 4 hours did not significantly reduced the population size of 40288.

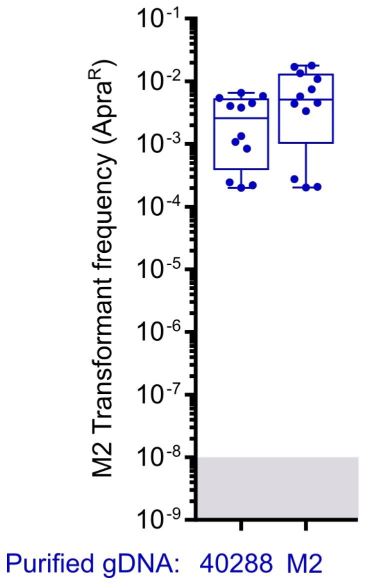


**Figure S2. Strain M2 of *A. nosocomialis* is equally transformed with DNA from itself and from the distantly-related strain 40288 of the *A. baumannii* species.**
Strain M2 was incubated under transformation conditions (>16 hours) with either genomic DNA extracted from a M2 Δ*comEC:aacC4* (ApraR) or from a 40288 Δc*omEC:aacC4* (ApraR) strain. Transformation frequency represent the CFU count on apramycin plate divided by the total CFU count on non-selective plates.

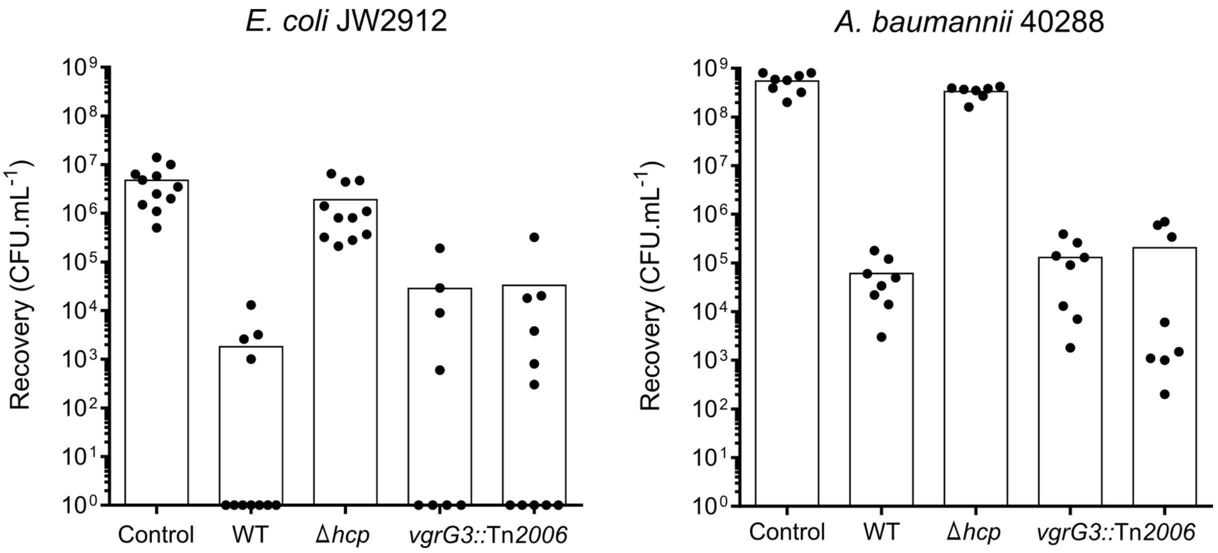


**Figure S3: T6SS-mediated killing assay of a VgrG3 impaired mutant of *A. nosocomialis* strain M2 in LB medium.**

Killing assays were performed as described in Figure S1. *Escherichia coli* strain JW2912 (Kan$^R$) and *A. baumannii* strain 40288 were both used as preys and were incubated either with the wild type M2 strain (WT), its *hcp* gene mutant derivative (Δ*hcp*) or with an imipenem resistant transformant carrying an insertion into the *vgrG3* gene (*vgrG3::*Tn*2006*, two transformants tested).

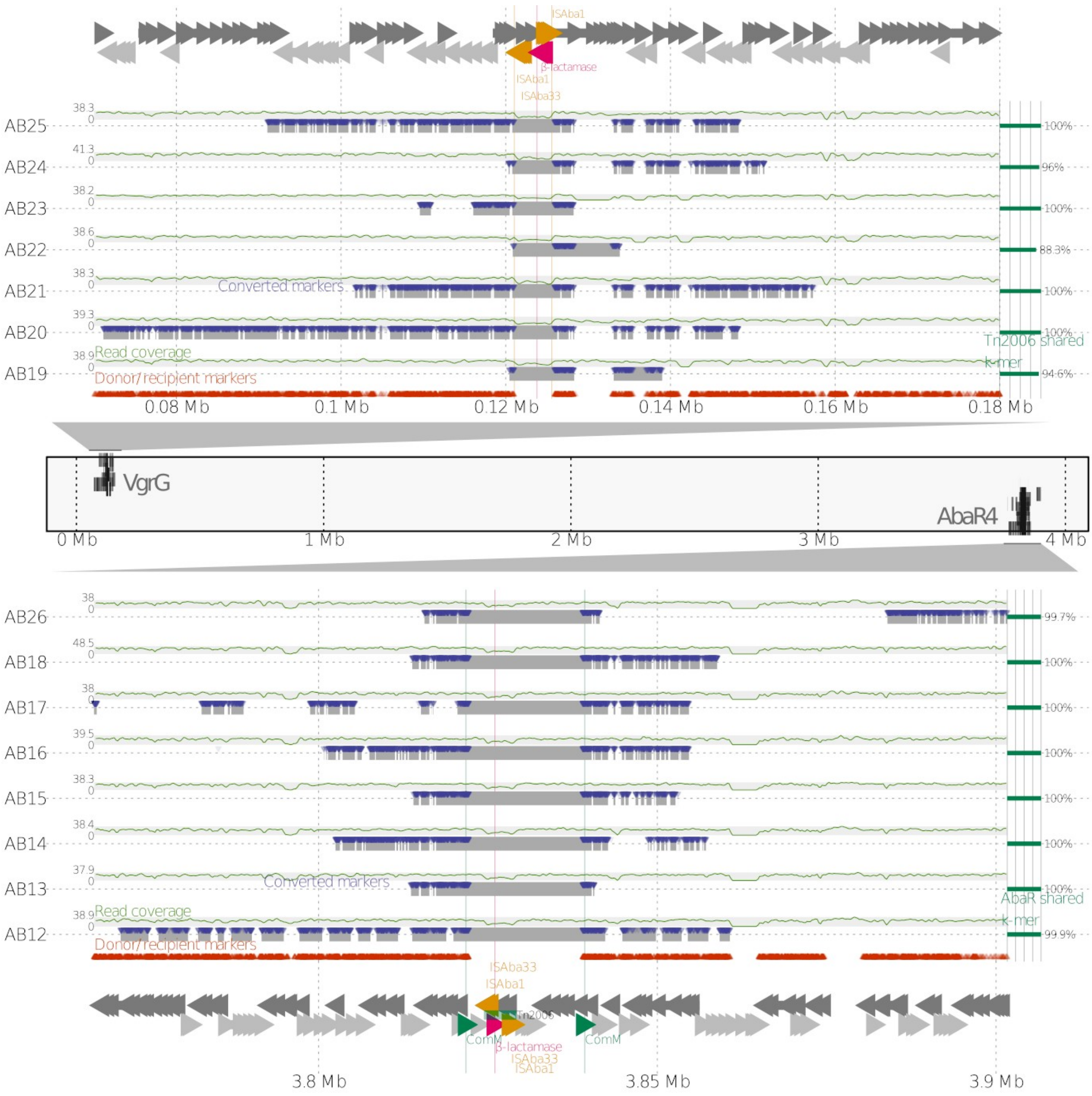


**Figure S4: Genomic analysis of the acquisition of imipenem resistance.**
Graphical representation of the chromosome of the 40288 strain (boxed area) and location of the acquired DNA fragments (light grey thick lines) by transformants of the *A. nosocomialis* strain M2 during mixed culture with 40288. Top and bottom panels represent close-up of the regions in which the *bla*$_{OXA-23}$ gene is acquired as part of the Tn*2006* element (top box) in the *vgrG3* gene and as part of the Tn*2006* element within the AbaR island (bottom box) inserted in the *comM* gene. Acquired regions were determined by sequencing the genomes of imipenem-resistant recombinants of M2 which had acquired *bla*$_{OXA-23}$ gene on a Novaseq instrument. Variant calling and identification of converted markers (SNPs of 40288 acquired by M2) were used to delineate the acquired regions (see Material and Methods). Thin green lines represent coverage and outermost right scale indicate the result of read screening corresponding to the AbaR sequence in recombinants. Orange triangle represent individual SNPs used to determine recombination tracts.

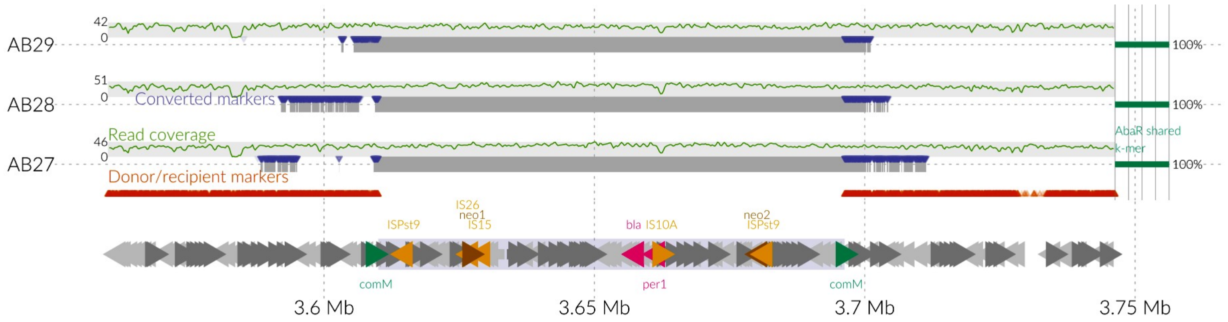


**Figure S5: Genomic analysis of the acquisition of the AbaR1 island**

Graphical representation of the acquired fragments and the AbaR1 island in the M2 transformants. Variant calling and identification of converted markers (SNPs of AYE acquired by M2) were used to delineate the acquired regions (see Material and Methods). Thin green lines represent coverage and outermost right scale indicate the result of read screening corresponding to the AbaR sequence in recombinants. Orange triangle represent individual SNPs used to determine recombination tracts.
